# Dorsal striatal dopamine induces fronto-cortical hypoactivity and implies reduced anxiety and compulsive behaviors in rats

**DOI:** 10.1101/2021.02.11.430770

**Authors:** Agata Casado-Sainz, Frederik Gudmundsen, Simone L. Baerentzen, Denise Lange, Annemette Ringsted, Isabel Martinez-Tajada, Siria Medina, Hedok Lee, Claus Svarer, Sune H. Keller, Martin Schain, Celia Kjaerby, Patrick M. Fisher, Paul Cumming, Mikael Palner

## Abstract

Dorsal striatal dopamine transmission engages the cortico-striato-thalamo-cortical (CSTC) circuit, which is implicated in many neuropsychiatric diseases, including obsessive-compulsive disorder (OCD). Yet it is unknown if dorsal striatal dopamine hyperactivity is the cause or consequence of changes elsewhere in the CSTC circuit. Classical pharmacological and neurotoxic manipulations of the CSTC and other brain circuits suffer from various drawbacks related to off-target effects and adaptive changes. Chemogenetics, on the other hand, enables a highly selective targeting of specific neuronal populations within a given circuit. In this study, we developed a chemogenetic method for selective activation of dopamine neurons in the substantia nigra, which innervating the rat dorsal striatum. We used this model to investigate effects of targeted dopamine activation on CSTC circuit function, especially in fronto-cortical regions. We found that chemogenetic activation of these neurons increased movement, as expected from dopamine release, rearings and time spend in center, while it also lowered self-grooming and increased prepulse inhibition in females. Remarkably, we observed reduced [^18^F]FDG metabolism in frontal cortex, following dopamine activation in the dorsal striatum, yet total glutamate levels-in this region were increased. A finding which may help explain the contradiction in some clinical studies of increased [^18^F]FDG metabolism and lower glutamate levels in diseases like OCD. Taken together, these results establish the importance of nigro-striatal dopamine transmission for modulating CSTC function, especially with respect to fronto-cortical activity, glutamate levels and behaviors related anxiety and compulsive actions.

**One Sentence Summary:** Dorsal striatum dopamine induce fronto-cortical hypoactivity and reduce compulsive behaviors in rats

## Introduction

Real progress in the development of therapeutic approaches for a range of neuropsychiatric disorder has been stalled for years. One reason is likely due to the lack of causal whole brain analysis of functional circuits, which are usually done for one region or projection at the time. We set out to discover how manipulation of one dopaminergic projection affects changes throughout the cortico-striato-thalamo-cortical (CSTC) circuit using *in vivo* whole-brain imaging and chemogenetics.

The CSTC circuit relays cortical activity, through the striatum, midbrain and thalamus back to the cortex. Cortical pyramidal neurons project to the dorsal striatum, where spiny projection neurons (SPN) then subserve an integration of direct (go) or indirect (no-go) output pathways that relay signals to the thalamus via the substantia nigra pars reticulata (SNr). The thalamus then communicates the processed signal back to frontal cortex to complete the circuit [2,6]. The activity of the CSTC circuit is modulated by nigro-striatal dopamine projections to the dorsal striatum. Changes in striatal dopaminergic activity have been linked to perturbations of cognitive functions in multiple neuropsychiatric or neurodegenerative diseases [1] such as obsessive compulsive disorder (OCD) [2] and schizophrenia [3]. While showing distinct phenomenologies, these conditions have overlapping behavioral phenotypes, including obsessions, compulsions, and sensory gating deficits, which are linked to altered dopaminergic transmission within the striatum [4,5].

Molecular imaging studies by positron emission tomography (PET) in people with OCD have shown changes in striatal dopamine receptor availability [5], metabolic hyperactivity especially in the orbito-frontal cortex (OFC), and increased metabolic connectivity from frontal cortex to the caudate [7–10], which is the equivalent of the medial dorsal striatum (mDS) in rodents [11]. MR spectroscopy studies of such individuals have shown relatively reduced total glutamate+glutamine (Glx) and N-acetylaspartate levels in fronto-cortical regions [12–14]. These clinical imaging studies suggest the presence of a phenotype of abnormal CSTC circuit activity in OCD, although it is difficult to establish causal relationships based on clinical imaging without direct targeting of specific neuronal pathways. As such, it is unclear if altered striatal dopamine function is a driver for or rather a consequence of fronto-cortical dysfunction in neuropsychiatric disease.

Classical neurochemical approaches to modulate activity within the rodent CSTC include neurochemical lesions [15,16], and focal stereotaxic drug injections [17]. These chronic methods, in varying degrees, bring unintended perturbations or adaptive changes of the nervous system. Chemogenetics, on the other hand, enables a highly selective targeting of specific neuronal populations within a given circuit. In this study, we applied a selective retrograde chemogenetic transduction of the medial dorsal striatum (mDS) dopamine projections arising from the medial substantia nigra pars compacta (mSNpc). Using this model, we studied the effects of selective nigro-striatal dopamine activation on behavioral, and whole-brain CSTC circuit function in the rodent brain. The combination of [^18^F]FDG PET and MR Spectroscopy gives valuable knowledge highly needed in order to interpret clinical neuroimaging findings in psychiatric disorders, and suggest that the combination of these using PET/MR technology in patients may increase our understanding of such disorders. We show that there is a causal relationship between increased dorsal striatum dopamine and fronto-cortical hypoactivity, yet unexpectedly also an increase in total glutamate levels. At the same time, dopamine activity in the dorsal striatum led to increased locomotion, increase exploratory and anxiolytic effects, and a change in behavioral measures related to CSTC circuit function and compulsive actions, lower self-grooming, lower startle response and increased prepulse-inhibition of the startle response in females. These results in otherwise behaviorally normal rats support the formulation of a novel hypothesis that potentiating dopaminergic activity specifically in mDS will reduce anxiety and compulsive-like behavioral symptoms which are related to hypercortical activity, like those present in disorders such as OCD.

## Materials and Methods

### Animals

Long Evans TH:Cre rats (LE Tg(TH:Cre 3.1)Deis) were purchased from the Rat Resource and Research Center (RRRC, University of Missouri). This strain carries a transgene containing a Cre recombinase inserted immediately before the start codon of the tyrosine hydroxylase gene (Witten et al. 2011). Both male (♂ = ▲) and female (♀ =▼) animal were used. Additional information of breeding and ethics is located in the supplementary materials.

### Statistics

Results are analyzed in GraphPad using a two-way ANOVA (behavioral experiments) or mixed effects model (PET and MRS) analysis with repeated measure where appropriate with adjustment for multiple comparisons. We report general effects of sex, treatment or vector injection with significance (P) and degrees of freedom (F), while mixed effects models are noted with t-value and DF. Individual group differences are noted only with respect to significance (P). Standard deviations (SD) are reported throughout the results and figures. Significance is reported as **** P<0.0001, *** P<0.001, ** P<0.01, * P<0.05.

### Retrograde chemogenetic transduction of dopaminergic projections in rats and mice

Adult TH:Cre rats (250 g) rats were anesthetized with 3% isoflurane (vol/vol) in oxygen and placed into a stereotactic frame (Kopf Instruments, Tujunga, CA, USA). Anesthesia was maintained with 1.5– 2% isoflurane; breathing, pain reflex, and body temperature was monitored throughout surgery. The scalp was shaved and disinfected with successive swabs soaked in 70% iodine, ethanol, and 0.1% lidocaine. An incision was then made down the midline of the scalp, the skull was exposed, and burr holes were drilled above the target regions. The DREADD (mDS-DREADD: AAV6-hSyn-DIO-mCherry-hM3D(Gq)-WPRE) (hM3Dq) and mCherry (mDS-mCherry: AAV6-hSyn-DIO-mCherry-WPRE) carrying viral vectors (described in detail in the supplementary materials) were injected bilaterally in the mDS at two locations in each hemisphere (2 μL per location) (AP +1.1 mm, ML ± 2.5 mm, DV −5 mm and −6 mm), via a 10 μl Nanofil syringe and 33GA beveled needle (World Precision Instruments, Sarasota, FL, USA) at an infusion rate of 150 nL/min. After infusion, the needle was left in place for ten min to allow for diffusion of the virus away from the needle tip before its slow withdrawal. Following injections, the scalp was sutured and the rats were allowed to regain consciousness. Carprofen (5 mg/kg) was administered subcutaneously as an analgesic before surgery and at 24 and 48 h after surgery. Following surgery, rats were single-housed for one week, and their weight, fur, eyes and overall movement were checked daily for signs of distress following the scheme of Roughan and Flecknell [18]. Animals were then returned to group housing for two weeks prior to further experiments.

### Immunohistochemistry

At the end of the experiments, rats were euthanized with an overdose of pentobarbital and immediately perfused transcardially with 150-200 ml of ice-cold phosphate-buffered saline (PBS, pH 7.4) followed by 100-150 ml of ice-cold 4% paraformaldehyde in PBS. Brains were removed and stored in 4% paraformaldehyde for 24 h at 4 °C, rinsed twice with PBS, and then cryoprotected in 30% sucrose/PBS. Brain sections 40 μm-thick obtained on a microtome were stored in cryoprotectant solution at −20 °C. Free-floating sections were immunostained for mCherry (Cat. #632543, TaKaRa Bio Europe) using the Avidin-Biotin Complex Peroxidase (ABC-P) method and for TH (Cat. #P21962, Invitrogen) using fluorescent antibodies as described in detail in the supplementary material.

### Locomotor activity and prepulse inhibition of the startle response

First, we assessed the individual startle responses of the rats as described in detail in the supplementary material. On the experimental day, locomotor activity and prepulse inhibition of the startle response were recorded from 43 animals in cohort 1 at baseline and again after CNO treatment (14 WT (7M / 7F), 15 mDS-DREADD (8M / 7F) and 14 mDS-mCherry rats (7M / 7F)). Locomotor activity was assessed in a custom-made open field maze consisting of an 80 x 80 cm arena enclosed by 65 cm opaque walls, with indirect, dim illumination (~10 Lux). On the test day, rats received an i.p. injection of CNO (0.5 mg/kg in 5% DSMO/Saline), and 20 min later were placed at the center of the open field and allowed to explore for 40 minutes. Behavior was recorded with a Basler ace2 Pro video camera with a 4.5-12.5 mm F1.2 lens positioned directly above the arena and analyzed using Ethovision software, with scoring of distance, velocity, center of field entries, rearing and grooming. The arena was cleaned with soap and water and then dried between each session. Immediately after the locomotion recordings, the rats were transferred to the startle chamber for sensorimotor gating assessment, details of which are described in the supplementary material.

### [^18^F]FDG PET scanning

The rats (n=14, (7 mDS-DREADD, 4F/3M), (7 mDS-mCherry, 4F/3M)) were fasted overnight before the PET scan. At the day of the experiment, they received an s.c. injection of 0.5 mg/kg CNO (0.5 mg/kg in 5% DSMO/Saline) or saline followed 20 min later by an i.p. injection of 14.1 ± 1.6 MBq FDG (from the clinical in-house production). The rats remained in their home cage for 45 minutes following the [^18^F]FDG injection. Following this uptake period, the rats were sedated with isoflurane (2–2.5% in oxygen) and placed in a homemade four rat insert in a Siemens HRRT (High Resolution Research Tomograph) scanner [19] for a 45-min list-mode emission scan. The rats were kept warm using an infrared lamp and monitored for respiration throughout the scan. PET images were cropped to brain-only images for each rat and manually co-registration to an-FDG-specific rat brain template. Individual scans were converted to SUV units and normalized to whole-brain uptake as described in detail in the supplementary material. The images were averaged per group/condition and the voxelwise difference was calculated between conditions. We undertook a regional analysis for each animal in Nucleus Acumbens (NAc), mDS, lateral Dorsal Striatum (lDS), Anterior Cingulate Cortex (ACC), mPFC, OFC, Thalamus (Thal), the midbrain including ventral tegmental area and substantia nigra (MB), Hypothalamus (Hyp) and Cerebellum (Cer), as described in detail in the supplementary material.

### Magnetic resonance spectroscopy

All MR scans were performed on a Bruker BioSpec 94/30 USR MRI system (9.4T, 30 cm bore, Bruker, Ettlingen, Germany) as previously described (29). In short, rats were anaesthetized with 2% isoflurane in oxygen, MRS was done sequentially in a 3 × 3 × 3 mm region in the right hemispheric mDS and then in a 3 × 2 × 3 mm region in mPFC. The spectra were acquired using a STEAM sequence with TE = 4 ms, TR = 4000 ms, 400 averages, eight dummy scans and 4096 points. VAPOR was used for water suppression. Metabolite concentration calculation was performed by LCModel using water unsuppressed signal as an internal reference.

## Results

### Selective chemogenetic targeting of nigro-striatal dopaminergic projections

We first confirmed the retrograde transduction of nigrostriatal dopamine neurons. Injection of Cre-dependent serotype 6 adeno associated viruses in the mDS of TH:Cre rats led to selective retrograde expression of the mCherry reporter gene in dopamine projection neurons arising from the mSNpc (Fig 1, A and B). Immunohistochemical identification of the mCherry reporter gene showed excellent selectivity towards the target neurons. Two different viral vectors were injected into the mDS, one carrying a Gq-coupled DREADD together with mCherry and the other only mCherry (Figure 1D). The staining intensity and spread from the injection site confirmed a highly reproducible target transduction in the mDS (Fig. S2A). There was no evident transduction of cell bodies resident in the mDS, nor of non-TH:Cre positive cells in the mSNpc (Fig. 1, E-J). Notably, the ventral tegmental area was devoid of mCherry-expressing somata (Fig. 1, E and I).

**Figure 1:**
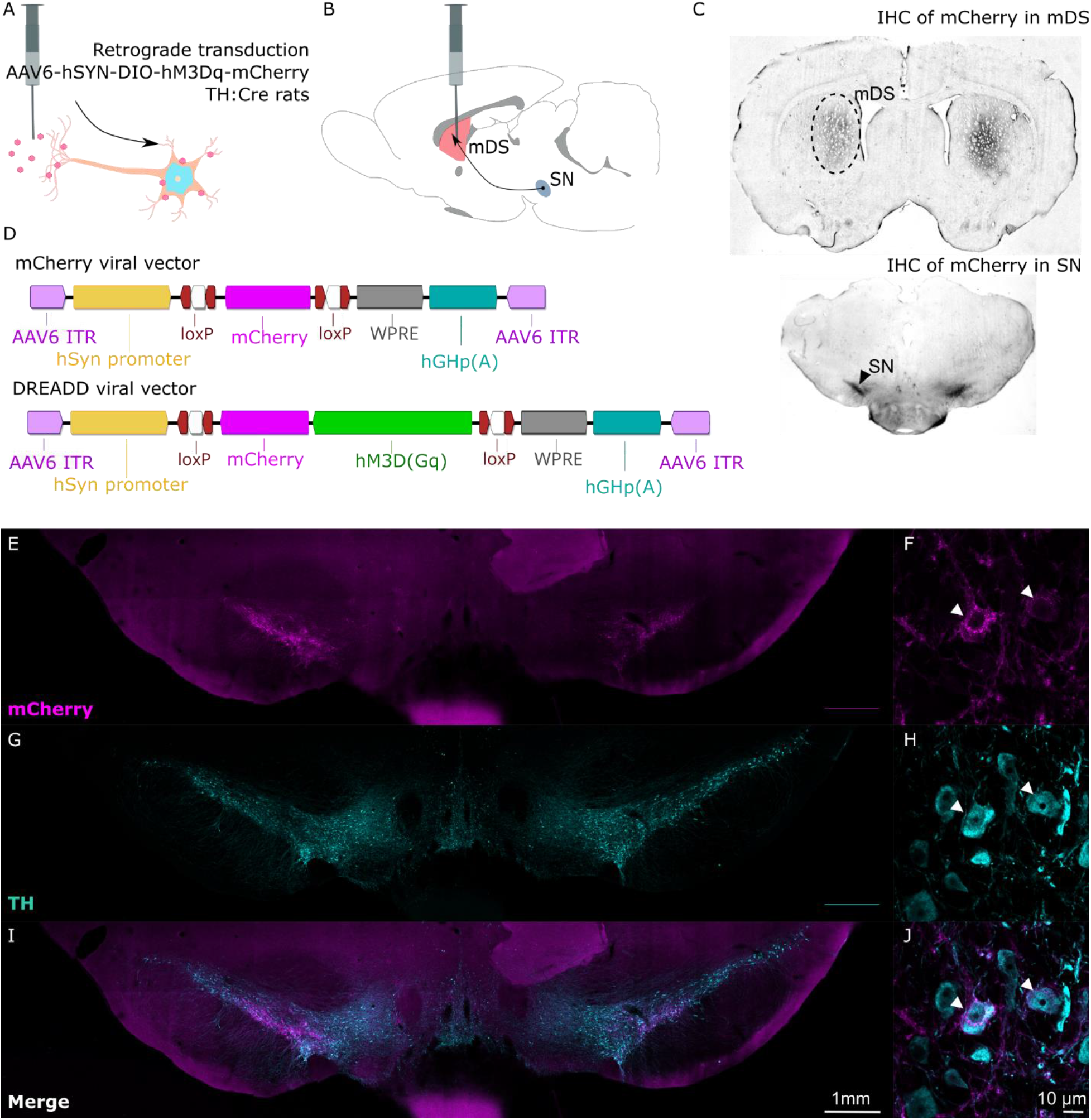
A,B) Retrograde transduction of Cre-positive tyrosine hydroxylase (TH) expressing neurons in the dorsomedial striatum. C) Immunohistochemical staining of mCherry in dorsomedial striatum and in substantia nigra. D) Viral vectors used in the vehicle (mDS-mCherry) and chemogenetic (mDS-DREADD) transduction groups. E,G,I) Fluorescent microscopy of mCherry (viral transducted) and TH-positive (dopamine) cells in the mesencephalon. F,H,J) Magnifications of individual transducted cells (two examples labelled with white arrows) in the SN.

### Nigro-striatal dopaminergic stimulation increases exploratory behaviors and lowers self-grooming

We then tested the efficacy of our transduction by locomotor activity. CNO significantly increased velocity in mDS-DREADD animals, with a maximal 90% increase occurring after 20 minutes of exploration, as compared with wildtype (WT) and mDS-mCherry animals (Fig. 2, A and B). The mDS-DREADD rats showed an increase in total distance travelled and total time spent in locomotion. WT rats had a 14% decrease in locomotion time following CNO treatment, which we attribute to a general habituation to the open field during the second exposure (Fig. 2E). Yet, the number of movement initiations (frequency) was similar in all groups (Fig. 2I). There was increased number of entries (+125%) and time spent (+119%) in the center of the open field for mDS-DREADD rats (P<0.0001, n=12) following CNO treatment (Fig. 2, F and J). Time spent rearing and number of rearing events were similarly increased by 66 and 99%, respectively (P<0.001, n=12) following CNO treatment in mDS-DREADD rats (Fig. 2, G and K). Self grooming time and bouts were likewise affected, but in the opposite direction, with significant reductions of −59% and −38%, respectively (P<0.001, n=12) following CNO treatment (Fig. 2, H and L). Similarly, for the second batch of animals used for the imaging experiments, there was a similar increase in total movement (P=0.003, n=9), center entries (P=0.005, n=9) and rearing (P=0.002, n=9) in mDS-DREADD animals following CNO treatment (Fig. S4, A-C). There were no main effects of transduction or sex on either behavior. Taken together, mDS-DREADD rats moved and explored more following CNO treatment compared to WT and mDS-mCherry rats, while spending less time self-grooming.

**Figure 2:**
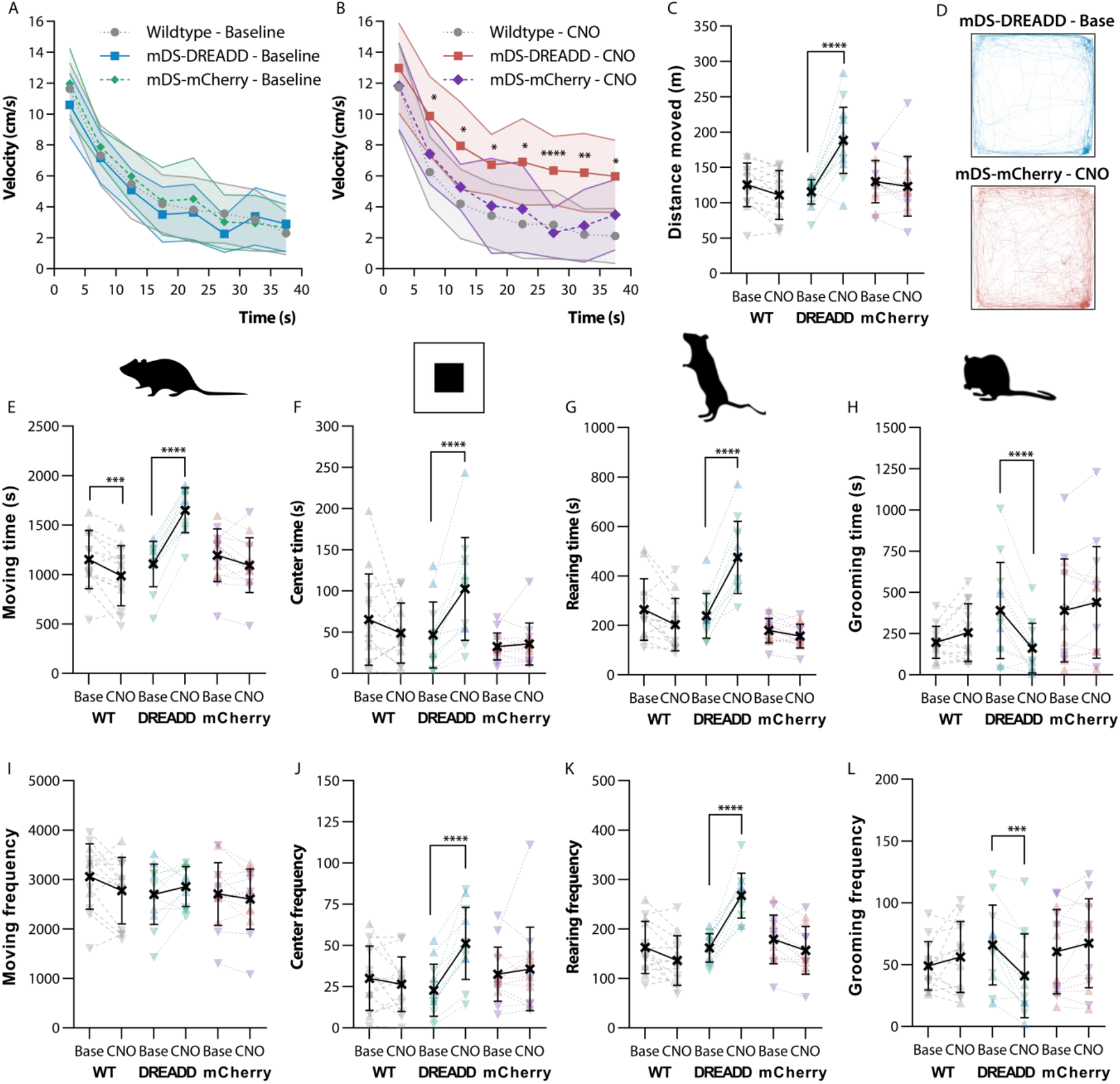
Open field behavior. A, B) Velocity over time in the open field. C) Total distance traveled over the whole-time course. D) Tracking of the median mDS-DREADD rat before and after CNO. E, F, G, H) Total time spent moving, in the center of the open field, rearing or grooming. I, J, K, L) Frequency of movement initiations, entries into the center square, rearings or grooming bouts. (♂ = ▲) (♀ =▼).

### Nigro-striatal dopaminergic stimulation affects prepulse inhibition of the acoustic startle response differently in males and females

Prepulse inhibition (PPI) of the acoustic startle response is modulated by CSTC circuit function and impaired in related neuropsychiatric disorders like OCD [20–22]. To assess PPI, we first tested basal acoustic startle response (ASR) to different intensity sound pulses. The ASR was clearly measurable after exceeding 100 dB amplitude and plateaued at 110 dB both in males and females, with a maximal ASR (as measured at 120 dB) of 59.1 ± 8.1% in males (n=21) and 49.6 ± 9.1% in females (n=27) (Fig. 3A). We chose 110 dB as the pulse intensity for subsequent testing of pre-pulse inhibition (PPI). There was a sex difference in the ASR to various volumes of sound pulses (P<0.001, F(1,46)=25.2), but no effect of transductions.

**Figure 3:**
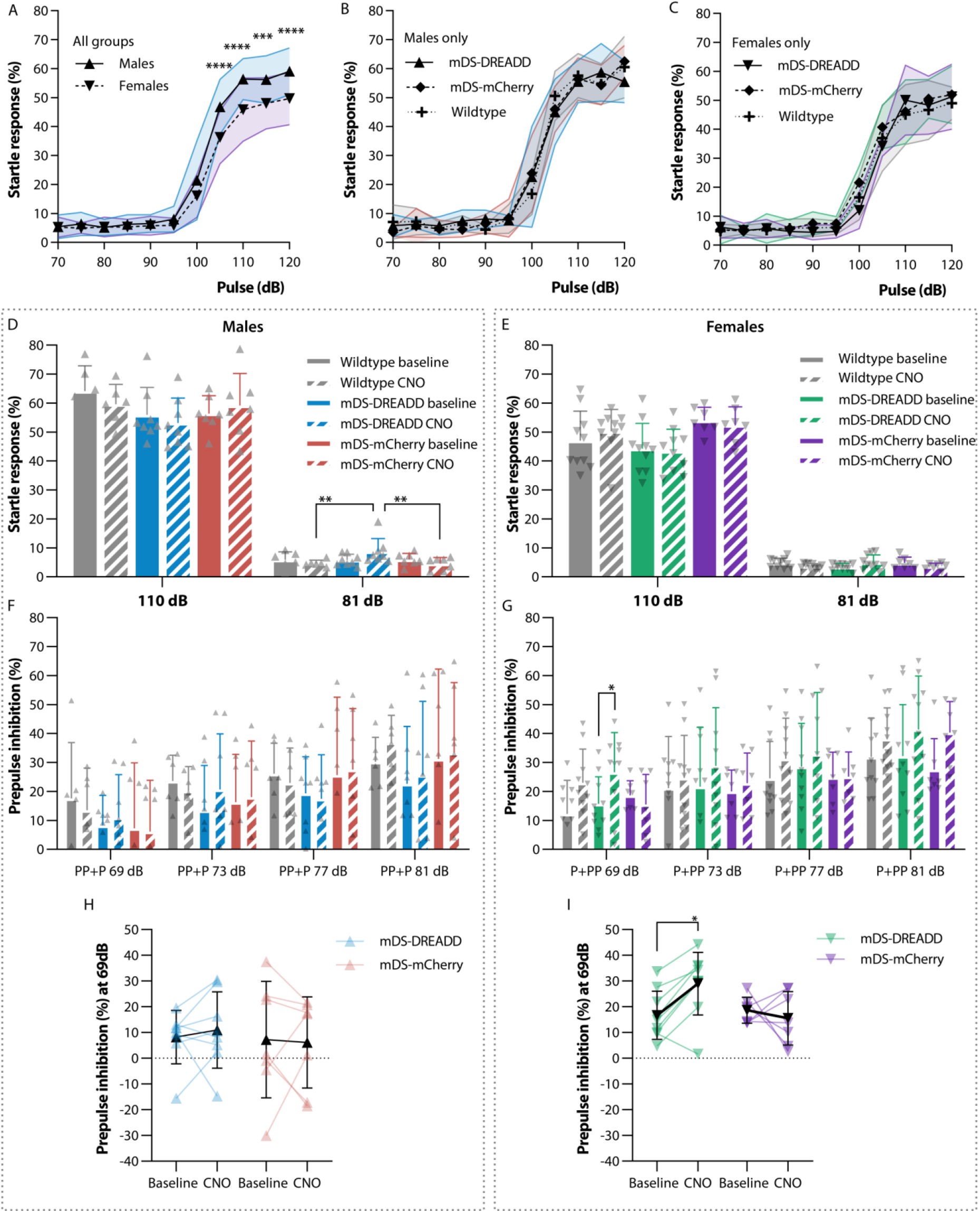
Acoustic startle response and prepulse inhibition. A, B, C) Startle response to various volumes of pulses. D, E) Startle response and effect of mDS-DREADD activation at high volume (110 dB) pulses and low volume (81 dB) prepulses. F, G) PPI as a result of prepulse intensities, negative PPI values are not shown in F and G to simplify the graph but were included in the statistical analysis. H, I) All PPI values at the lowest prepulse intensity (69 dB) pre and post CNO treatment to activate the nigro-striatal dopaminergic pathway. (♂ = ▲) (♀ =▼).

We then tested startle response (SR) to a 110 dB sound pulse with or without different intensity prepulses (all pulses are shown in Fig. S5A and D) and short-term habituation (STH) (Fig. S5B and E). There was no effect of transductions or treatment with CNO on the SR (Fig. 3, D and E), but there was an overall effect of sex. SR was greater in males (58.1 ± 3.8%, n=21) than in females (48.6 ± 4.3%, n=27) (P<0.001, F(1,84)=30.64), as expected from the initial ASR test (Fig. 3A). There were no differences in mean STH between WT and transduction groups or between baseline and CNO treatment. However, the mean STH was significant in all female groups combined (19.7 ± 6.2%; n=27) (Fig. S5B and E) (P=0.008, F(1,86)=7.3).

In general, SRs decreased in proportion to the amplitude of the prepulse, as expected (Fig. S5A and D). There were no general effects of transduction or CNO treatment on SR in either sex. Yet, a reduced SR following CNO treatment in mDS-DREADD females was just shy of statistical significance (Ave. Diff. = 3.3 ± 1.5 %, p=0.0501 (F(1,9)=5.11)). We next assessed the latency from startle pulse to startle onset and maximum peak of startle from the SR data above (Fig. S5C and F). There were no differences in latency with prepulse only and no stimuli (Fig. S5H). Baseline PPI following the 69 dB prepulse was higher in female than male rats (Fig. S5G) (P=0.006, F(1,84)=7.86). Transduction did not affect baseline PPI (Fig. 3F and G, and Fig. S5G). There was a general 24% increase in mDS-DREADD female rats for all pre-pulses amplitudes (P=0.022, F(1,9)=7.6), and a 58% increase in female mDS-DREADD rats at the 69 dB prepulse condition upon CNO treatment (Fig. 3G and I) (P=0.018, n=10).

### Nigro-striatal dopaminergic stimulation affects metabolic activity the medial prefrontal cortex and orbitofrontal cortex

Next, we wanted to know how dorsal striatal dopamine effected clinical neuroimaging measurements often used to study neuropsychiatric disorders. We used a whole-brain [^18^F]FDG PET approach to assess the glucose metabolism in the whole CSTC circuit at once. An image analysis of voxel intensity showed a relative lower uptake of [^18^F]FDG in cortical regions and higher uptake in cerebellar regions in mDS-DREADD rats after CNO treatment (Figure 4A). On a CSTC regional level, there were significant reductions in [^18^F]FDG uptake in the mPFC (−9%, P = 0.002, t = 4.06, DF = 60, N = 7) and OFC (−8%, P = 0.004, t = 3.73, DF = 49, N = 8) following CNO treatment in mDS-DREADD rats. There were no effects on [^18^F]FDG uptake in other CSTC regions of mDS-DREADD, nor in mDS-mCherry rats (Fig. 4B), and no effects of sex or transduction.

**Figure 4:**
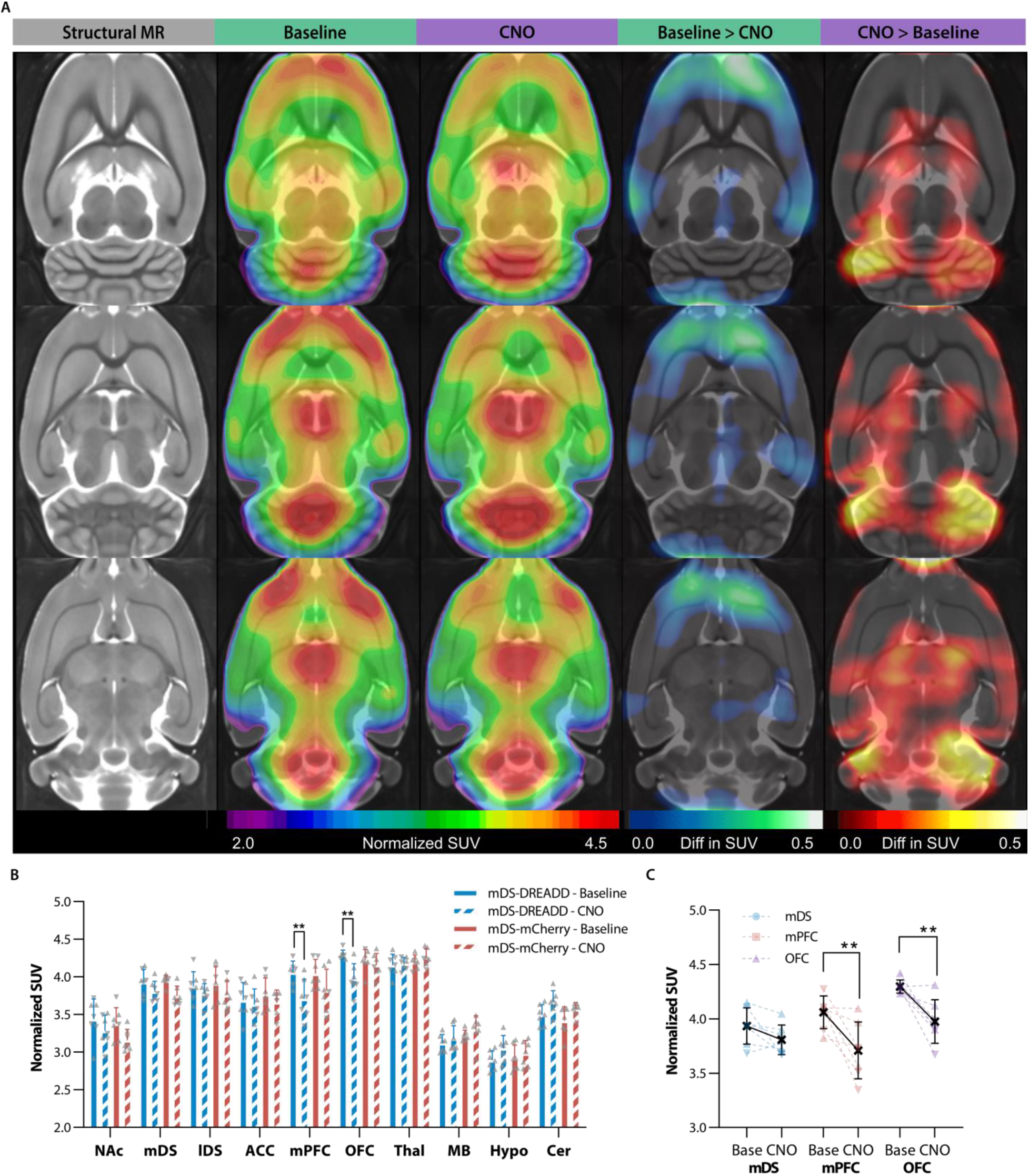
[^18^F]FDG images. A) [^18^F]FDG uptake in mDS-DREADD animal at baseline and after chemogenetic (CNO) stimulation (rainbow color), lower metabolic activity (blue color), higher metabolic activity (red color). B) Regional standard uptake values at all conditions and C) significant changes and individual animals in mDS, mPFC and OFC. (♂ = ▲) (♀ =▼). Nucleus Acumbens (NAc), medial Dorsal Striatum (mDS), lateral Dorsal Striatum (lDS), Anterior Cingulate Cortex (ACC), medial Prefrontal Cortex (mPFC), Orbito Frontal Cortex (OFC), Thalamus (Thal), the midbrain including ventral tegmental area and substantia nigra (MB), Hypthalamus (Hyp) and Cerebellum (Cer).

### Nigro-striatal dopaminergic stimulation increase glutamate and N-acetylaspartylglutamic acid in medial prefrontal cortex and glutamate levels in dorsal striatum

Based on the [^18^F]FDG PET findings (Fig. 4B), we chose two regions (mPFC and DS) for MR spectroscopy (MRS) measurements of glutamate, glutamine, N-aspartate (NAA) and N-acetylaspartylglutamic acid (NAAG) *in vivo.* There were significant increases in glutamate (P < 0.0001, t = 5.73, DF = 70), total NAA+NAAG (P = 0.006, t = 3.43, DF = 70) and total glutamate + glutamine (P = 0.001, t = 3.95, DF = 70) levels in the mPFC following CNO treatment in mDS-DREADD rats (Fig. 5, D and E). Furthermore, significant increases in glutamate (P < 0.0001, t = 5.67, DF = 70) and total glutamate + glutamine (P < 0.0001, t = 5.54, DF = 70) levels were found in the mDS. We found no effect of sex in any group, nor for viral transductions in baseline conditions. Contrary to our previous findings [23], CNO treatment in mDS-mCherry animals did provoke significant reductions in glutamate (P = 0.0002, t = 4.69, DF = 70) and total glutamate + glutamine (P < 0.0001, t = 6.01, DF = 70) levels (Fig. 6, F and G) in the mDS.

**Figure 5:**
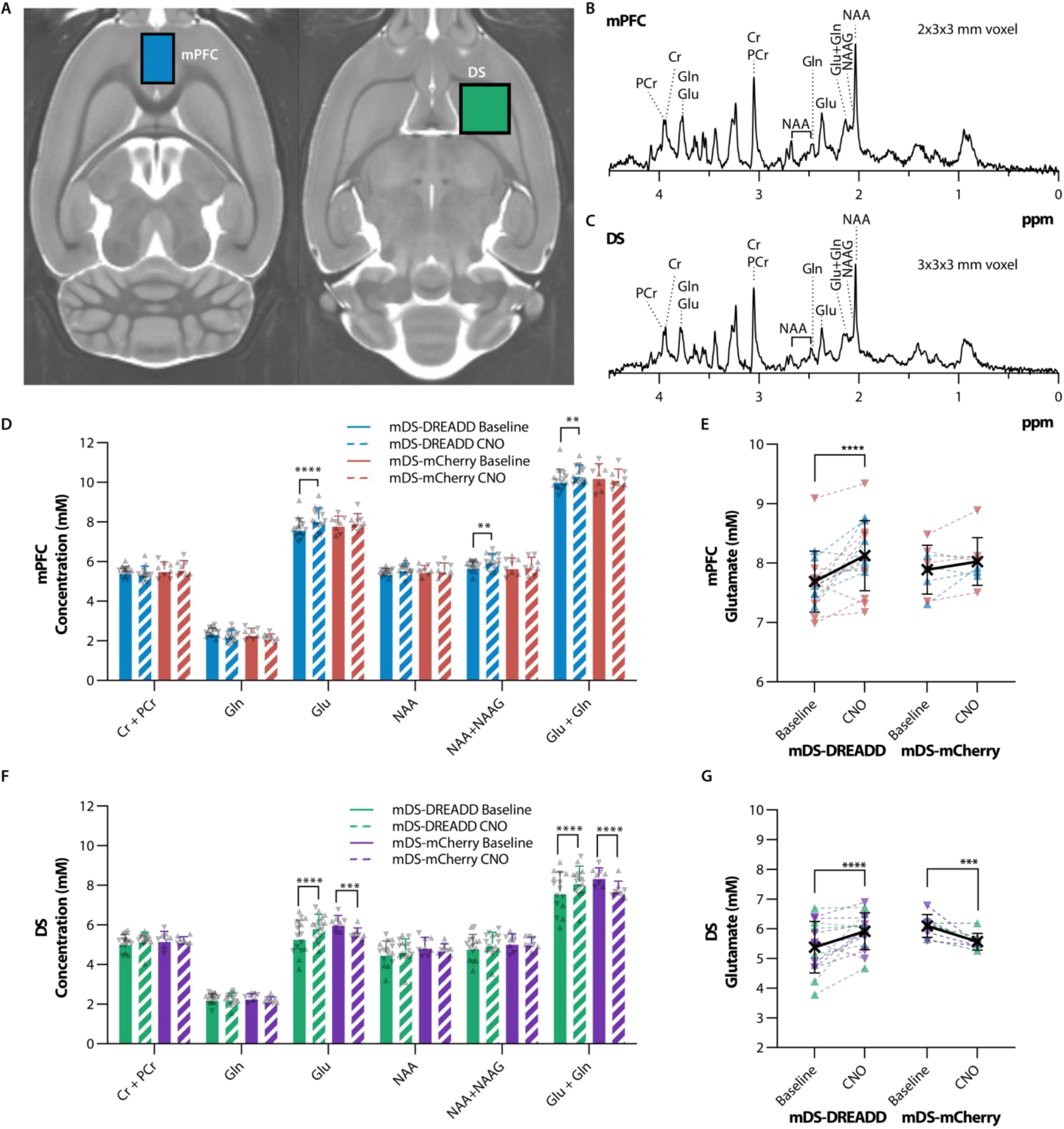
MR spectroscopy of neurochemicals in mDS-DREADD and mDS-mCherry animals. A) Voxel placement in mPFC (blue voxel) and DS (green voxel). B,C) Sample spectra from the mPFC and DS. D,F) Neurochemical concentrations in the mPFC in all conditions, E,G) significant changes in glutamate on the individual level. (♂ = ▲) (♀ =▼). Medial Prefrontal Cortex (mPFC), Dorsal Striatum (DS), Creatine (Cr), Phosphocreatine (PCr), Glutamine (Gln), Glutamate (Glu), N-acetyl-aspartate (NAA) and N-acetyl-aspartatylglutamate (NAAG).

## Discussion

We present a novel chemogenetic procedure for selective bilateral activation of nigro-striatal dopamine projections innervating the rat mDS, which enabled us to test effects of dopaminergic modulation of the CSTC circuit. In this animal model, we tested the specific hypothesis that selective dopamine activation in the rat mDS would lower compulsive-related behaviors and reduce cerebral metabolism in the frontal cortex. The least invasive, yet regional selective, procedure for rodent studies may be chemogenetics, in which transduction of Designer Receptors Exclusively Activated by Designer Drugs (DREADD) renders target neurons sensitive to G-protein coupled activation/inactivation following injection of an exogenous ligand, such as clozapine N-oxide (CNO) [24]. By enabling highly selective and reversible activation of specific neuronal subtypes and pathways, chemogenetics presents a powerful approach for establishing causal relationships.

### Nigro-striatal dopaminergic stimulation increases exploratory behaviors and lowers self-grooming

We targeted subdivisions of striatal dopaminergic innervations by using TH:Cre rats [25] treated with retrograde neuronal transduction by the AAV6 serotype, as previously established in rats [26], mice [27], and non-human primates [28]. We found that CNO increased lomomotor activity in all mDS-DREADD rats. *In vivo* stimulation of hM3Dq DREADDs are well known to induce phosphatidylinositol hydrolysis [24], c-fos activation [29–31], dopamine release [32], increased dopamine neuron excitability with a burst firing pattern [30,31,33,34] and ultimately increased locomotion as a behavioral read-out [30,35–39], which support our findings. We also observed an increase in exploratory behaviors, entries into center field, and rearing in all mDS-DREADD rats during chemogenetic activation. Such exploratory behaviors typically increase following treatment with anxiolytic drugs [40,41], and decrease following stress [41,42], often in a manner independent of hyperlocomotion, thus suggesting anxiolytic effects of the present perturbation of the CSTC circuit.

Self-grooming is an innate rodent behaviour characterized by a sequential pattern of movements [43], which serves as a useful proxy for compulsive behaviour [22,44,45]. Grooming time and frequency are under control by the efferent activity of striatal SPNs. Whereas activation of D_1_ receptors on SPNs of the direct (“go”) pathway increase grooming, activation of D_2_ receptors on the indirect (“no-go”) SPNs decrease grooming [43]. Our mDS-DREADD rats showed a significant decrease of grooming frequency and duration during chemogenetic activation. This may reflect preferential activation of the (D2-expressing) indirect pathway SPNs, although we cannot rule out the possibility of secondary effect due to hyperlocomotion and increased exploration. The preferential activation of the indirect pathway in our mDS-DREADD rats is supported by a recent report that the dopamine fibres, which was mapped by retrograde tracing, innervating the mDS predominantly target the “indirect” D_2_-positive SPNs [46]. Furthermore, a study by Ahmari et al, showed that hyperactivation of the OFC to DS pathway, increased self-grooming in rats, which support our findings where lower OFC activity lead to lower self-grooming [45].

### Nigro-striatal dopaminergic stimulation affects prepulse inhibition of the acoustic startle response differently in males and females

PPI is an autonomous behavioral response, which is highly modulated by components of the CSTC circuit, in particular the PFC [47–50], striatum [51], and the SN [52]. Disruption of normal PPI is evident in people with OCD [21,53]. Highly conserved across species, the PPI response serves as a useful translational construct in brain research. While some dopamine agonists disrupt PPI when given systemically, the relative contributions of D_1_-and D_2/3_ receptor pathways in this effect are debated [54–57]. Thus, converging lines of clinical and preclinical evidence link the CSTC circuit, dopamine, and PPI. The expression of PPI differs between rodent strains [58,59] and sex [60–62]. Indeed, we found that mDS-DREADD female rats had increased PPI at low prepulse amplitude during chemogenetic activation, but males showed no such effect. Previous work showed that male rats with SN lesions [52] as well as male mice with striatal lesions [63] displayed disruption of PPI, but there was no corresponding investigation of female rats in those studies. In another investigation, deep brain stimulation (DBS) of the NAc disrupted PPI in male WT rats, whereas the same DBS rescued disrupted PPI in a male rat model of schizophrenia [49], which implies context-dependence of the behavior consequences of stimulated dopamine release.

A limitation to this study lies in our use of CNO to activate DREADDs. CNO metabolizes *in vivo* to clozapine, which is presumably the main agonist on DREADDs [23,64–66]. While clozapine is an antagonist at endogenous dopamine D_2/3_ and serotonin 5-HT_2A_ receptors, it has selectivity for DREADDs when administered at the present low dose. Indeed, we did not observe any behavioural effects of CNO/clozapine in control rats in this study, nor had we seen any occupancy at D_2/3_ or 5-HT_2A_ receptors in an earlier *in vivo* binding experiment [23]. Furthermore, clozapine treatment can restore disrupted PPI [56,59,67], but neither we nor others [65] have measured any intrinsic effects of CNO on PPI in WT or control animals at the dose used, which also argues for selective activation of DREADDs.

### Nigro-striatal dopaminergic stimulation affects metabolic activity the medial prefrontal cortex and orbitofrontal cortex

Metabolic mapping of [^18^F]FDG uptake by PET can identify perturbation of neuronal circuits related to behavior, chemical stimulation, or chemogenetic-induced circuit modulation [49,50,68–71]. We found that chemogenetic activation in mDS-DREADD rats elicited hypometabolism in fronto-cortical areas (Fig 4A and B). This confirms the behavioral findings described above suggesting that mDS dopamine activation controls fronto-cortical neuronal activity though the CSTC circuit. Serveas et al. showed that acute treatment of rats with the dopamine D_2/3_ receptor agonist quinpirole provoked a global increase in [^18^F]FDG whole-brain uptake, yet a relatively lesser increase in NAc, mDS, and the fronto-cortical regions, i.e. mPFC, ACC and OFC [71], suggesting a common mechanism to the present DREADD study. Globally increased [^18^F]FDG uptake might hide regional differences in metabolism, emphasizing the importance of whole-brain normalization between baseline and test conditions [72]. Our use of the human HRRT PET scanner, which has lesser spatial resolution (2 mm) compared with contemporary dedicated small animal scanners (1 mm), limits our ability to detect small focal changes in metabolism. We accommodate the resolution issue by using relatively large bilateral VOIs.

### Nigro-striatal dopaminergic stimulation increase glutamate and N-acetylaspartylglutamic acid in medial prefrontal cortex and glutamate levels in dorsal striatum

Our MR spectroscopy results showed increases in glutamate and NAAG in the mPFC, and increased glutamate in the mDS during chemogenetic activation, which may seem at odds with the PET finding of lower [^18^F]FDG uptake in frontal cortex. While the exact relationship between [^18^F]FDG uptake and total glutamate levels is unknown, we suppose that glutamate is stored in the terminals of glutamatergic afferents in the mPFC, which are presumably thalamic or cortical inputs through the CSTC circuit, while NAAG is mainly located in interneurons [73]. Once released into the synapse, glutamate is taken up by astroglial cells and converted into glutamine, whereas NAAG is metabolized to NAA and glutamate [74,75]. In this scenario, the increased glutamate and NAAG levels measured by MR spectroscopy during chemogenetic activation may thus reflect elevated vesicular concentrations in thalamic or cortical afferent terminals, due to indirect inhibition via mDS dopaminergic activation of the indirect (no-go) pathway. NAAG signaling is not completely understood, but some evidence points towards it having an inhibitory effect on GABA release, such that lower NAAG release might result in net inhibition of cortico-striatal projections [76]. Such an inhibition could account for the lower cortical [^18^F]FDG uptake and consequently lower glutamatergic activity in the DS, resulting in increased vesicular glutamate content in DS. Furthermore, we suppose that mDS-DREADD stimulation also stimulates dopamine neurons that co-release glutamate [77,78], which in turn might favor the local astroglial conversion of glutamine to glutamate [74,75].

As noted above, there is no general model to connect glutamate levels measured with MRS to [^18^F]FDG uptake. One study reported no correlation between these two markers in WT mice, although there was a clear inverse relationship in mGluR5 KO mice [79]. That finding resembles present observations in the mDS-DREADD group, suggesting a similar perturbation of the coupling between glutamate levels and energy metabolism. We did observe a change in DS glutamate levels in the mCherry animals, as in our previous study, albeit at higher doses of CNO [23]. Thus, it remains possible that CNO metabolism to clozapine may indeed contribute to the cerebrometabolic changes reported in this study, albeit in the opposite direction; our results may therefore underestimate the effect of mDS dopamine activity on local glutamate levels

### Translational relevance

Changes in striatal dopamine and prefrontal cortical glutamate levels are held to be central etiological factors for neuropsychiatric disorders that involve the CSTC circuit, such as OCD [13] and Tourette’s Syndrome [80]. In a recent PET study with DBS of the ventral striatum in people with treatment-resistant OCD, increased dopamine release correlated with a decline in OCD related symptoms [81]. Furthermore Nordstrom et al. found that systemic administration of the D_2/3_ receptor agonist bromocriptine suppressed “Tics” in a transgenic mouse model of Tourette’s Syndrome [82], supporting an earlier clinical report that bromocriptine may be helpful in some OCD patients [83]. Our results are a step towards disentangling the functional and metabolic effects of specific elements in the CSTC circuit and support the proposition that mDS might be a relevant DBS target in neuropsychiatric diseases involving compulsive or impulsive behaviors.

### Conclusion

We found that chemogenetic activation of dopamine neurons projecting to mDS potentiated rat exploratory behavior, lowered self-grooming, and enhanced prepulse inhibition of the startle response. These behavioral changes occurred in association with reduced metabolic activity in frontal cortex as measured by [^18^F]FDG PET and increased glutamate and NAAG levels in the mPFC as measured by MRS. Our study thus revealed that dopamine signaling in mDS exerted modulatory control of fronto-cortical metabolic activity and the orchestration of behavior related to CSTC circuit function. These results support our hypothesis that dopamine release in the mDS can be a driver for reduced cortical metabolism in distal parts of the CSTC circuit and suggest that dopamine activation in the mDS reduce anxiety and compulsive behaviors.

## Acknowledgments

Structural reference MR image is generously provided by Kristian Nygaard Mortensen, Center for Translational Neuromedicine, University of Copenhagen. Technical expertise regarding acoustic startle response and prepulse inhibition was generously provided by Kim Fejgin at H. Lundbeck. Radiochemical support was generously provided by Matthias M. Herth and Vladimir Shalgunov, Department of Drug Design and Pharmacology, University of Copenhagen. Statistical discussions with Professor Todd Ogden, Department of Biostatistics, Columbia University, NY. Professor, DM Gitte M. Knudsen, head of the Neurobiology Research Unit, Copenhagen University Hospital, offered her support and guidance for students and researchers throughout the project. Funding for this project was provided to Mikael Palner by the Lundbeck Foundation (R192-2015-1591 and R194-2015-1589), Augustinus Foundation (18-3746 and 17-1982, Independent Research Fund Denmark (5053-00036B), Savværksejer Jeppe Juhls og Hustrus Ovita Juhls Mindelegat and Købmand i Odense Johann og Hanne Weimann født Seedorffs Legat.

## Supplementary Materials

Study Design

Chemicals

Viral Vectors

Software

Breeding and Phenotypic Assessment in Animals

Immunohistochemistry details

Acoustic startle response

Prepulse inhibition of the acoustic startle response

[^18^F]FDG PET scanning

### Supplementary Figures

Fig. S1. Experimental setup.

Fig. S2. Quantification of transduction in rats.

Fig. S3: Second batch of locomotor experiments in animals for imaging experiments.

Fig. S4: Additional acoustic startle data.

Fig. S5: A selected [^18^F]FDG-PET rat brain image with all atlas VOIs

