## Supplementary Material for "Dorsal striatal dopamine induces fronto-cortical hypoactivity and implies reduced anxiety and compulsive behaviors in rats" (from another version of this preprint: version 1, 2021-02-11)

### Title: Nigro-striatal dopamine activation lowers behavioral and neuronal phenotypes associated with obsessive-compulsive disorder

**Authors:** Agata Casado-Sainz[1]†, Frederik Gudmundsen[1]†, Simone L. Baerentzen[1], Denise Lange[2], Annemette Ringsted[1], Isabel Martinez-Tajada[1], Siria Medina[1], Hedok Lee[3], Claus Svarer[1], Sune H. Keller[4], Martin Schain[1], Celia Kjaerby[5], Patrick M. Fisher[1], Paul Cumming[6,7], Mikael Palner[1]*

#### Supplementary Materials and Methods

##### *Chemicals*

Clozapine N-oxide (CNO) purchased from Cayman Chemical Company (Cat. #16882) was diluted to 10 mg/ml in dimethyl sulfoxide (DMSO, 99% Merck) and stored at -20 ºC. Before use, it was further diluted 20-fold in normal saline and administered intraperitoneally at a dose of 0.5 mg/kg.

##### *Viral vectors*

The DREADD (hM3Dq-containing) adeno-associated virus (AAV) – rAAV6-hSYN-DIO-hM3Dq-mCherry – was produced by the Vector Core at the University of North Carolina at Chapel Hill (UNC Vector Core). The control (hM3Dq-negative) virus, – rAAV6-hSYN-DIO-mCherry – was produced by the Viral Vector Facility at the University of Zürich (UZH).

##### *Software*

We used the following commercial programs for data analysis and plotting: Matlab vR2018b (Mathworks), PMOD v3.7 (PMOD Technologies LLC), RStudio v1.2.1335 *(95)* (RStudio), FSL 5.0.11 (FMRIB Software Library), Prism v8 (GraphPad). EthoVision v.11.5 (Noldus Information Technologies), LC Model (Provencher).

##### *Breeding and Phenotypic Assessment in Animals*

TH:Cre positive males were crossed with wildtype females and outbred, resulting in hemizygous TH:Cre positive or wild-type pups. The breeding was conducted at the Department of Experimental Medicine (AEM, University of Copenhagen). The rats were ear-marked and transferred to the rodent facility at Rigshospitalet to habituate for at least one week before surgery. Animals were genotyped using the REDExtract-N-Amp™ Tissue PCR Kit (Cat. #XNAT, Sigma) and Cre recombinase oligonucleotides (Eurofins): forward 5'-GCG GTC TGG CAG TAA AAA CTA TC-3' and reverse: 5'-GTG AAA CAG CAT TGC TGT CAC TT-3'. The rats were housed in same sex groups of 2-3 animals per cage with 12 h light cycle beginning at 0600 and ending at 1800. All experiments were conducted during the light phase. Females were fed *ad libitum* (1319 Avlsfoder, Rotte/Mus, Brogaarden), but males were diet-restricted after their eighth week of age to 20 g of food pellets per day to keep their body weight below 500 g, as required for scanning. We conducted all procedures in accordance with the FELASA guidelines and with approval from The Danish Animal Experiments Inspectorate (license number: 2016-15-0201-01031). GMO procedures were approved by the Danish Working Environment Authority (Lab ID 229 806).

#### Supplementary Figures

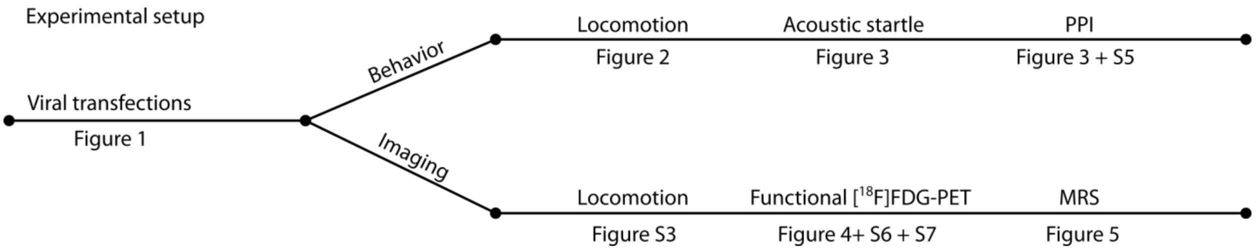


Figure S1: Experimental setup

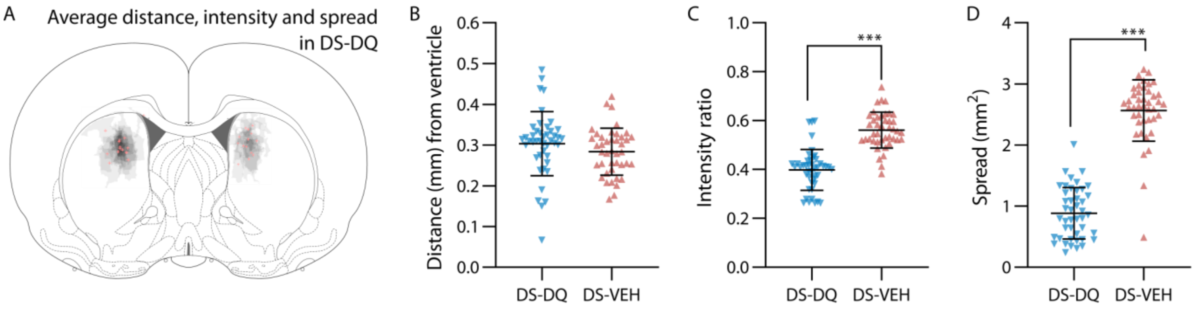


Figure S2: Quantification of transfection in rats, A) Visual representation of the average distance, intensity and spread of DS-DQ viral transfections. B,C,D) Quantified average distance, intensity and spread of DS-DQ viral transfections in relation to the smaller vector used in DS-VEH.

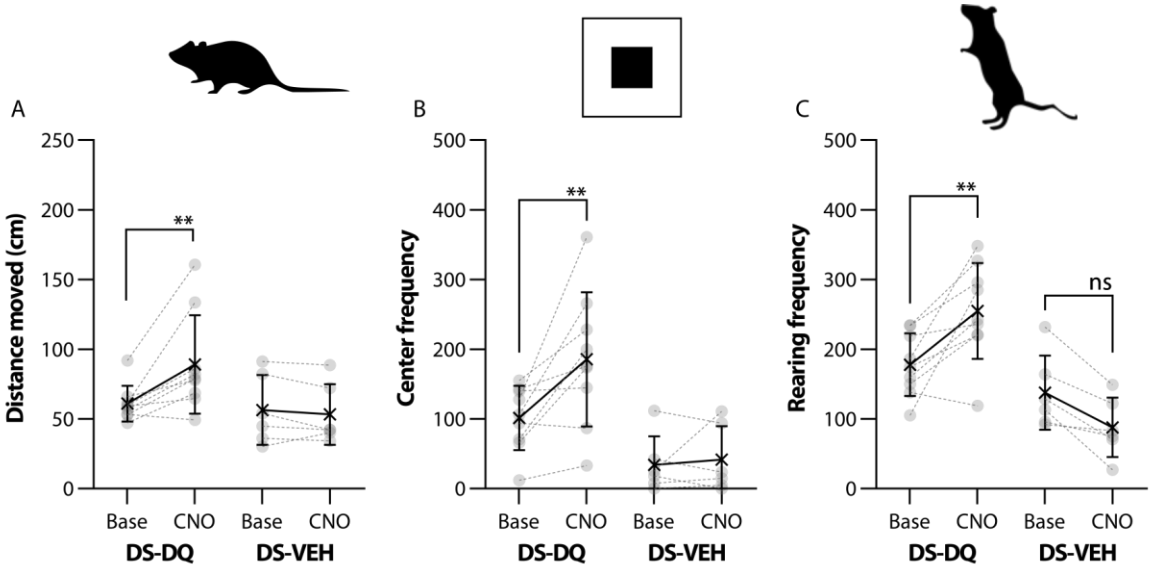


Figure S3: Second batch of locomotor experiments in animals for imaging experiments. A: 46% increase, from 610 cm to 890 cm, P = 0.0031. B: 82% increase, from 102 cm to 186 cm, P = 0.0053. C: 43% increase, from 178 cm to 255 cm, P = 0.0017

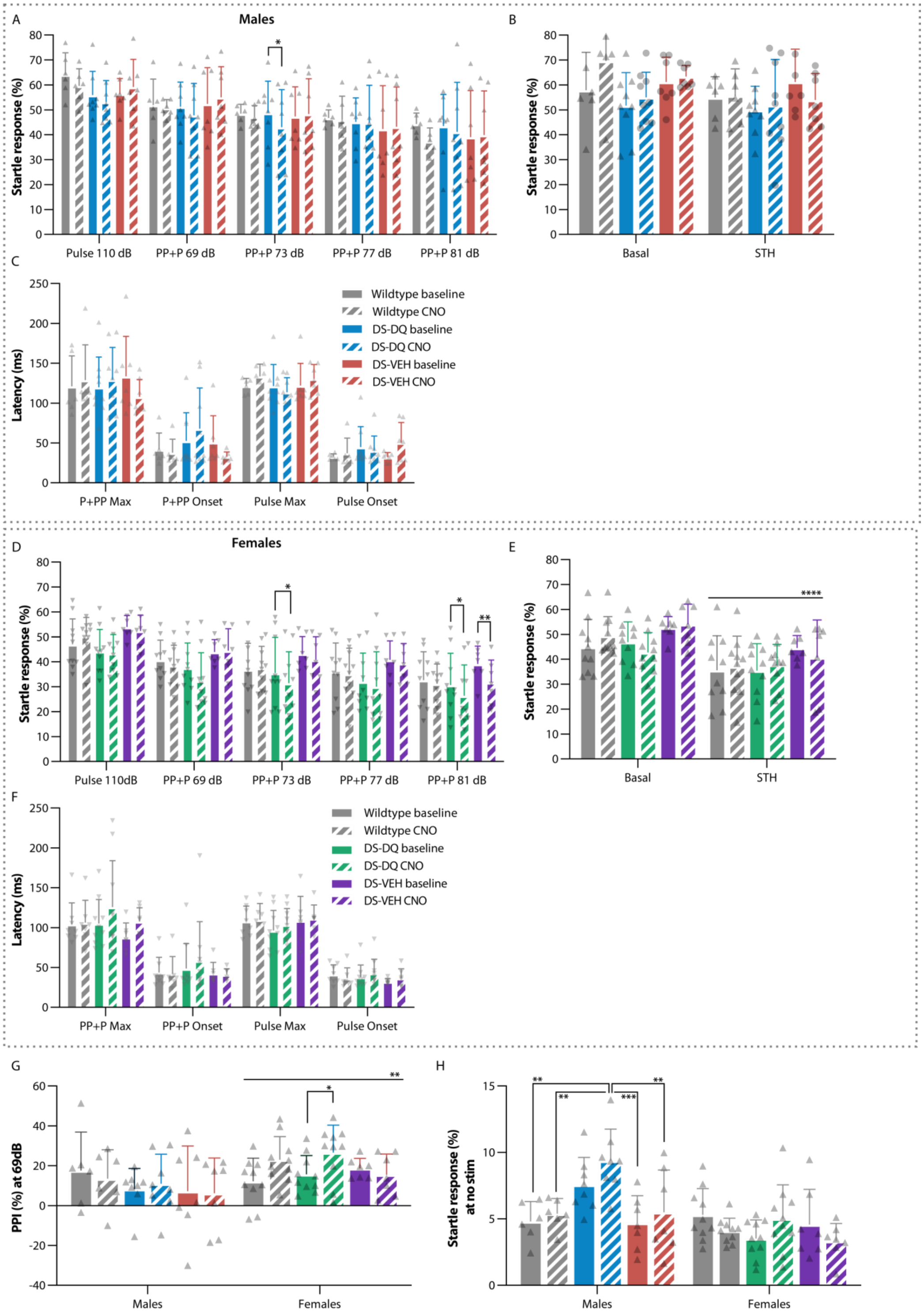


Figure S4: Additional acoustic startle data in males (top) and females (middle). A,D) Startle response following pulse and pulse + prepulse (PP+P) at each dB. B,E) Short term habituation within a startle sessions. C,F) Latency from pulse to startle. G) Sex comparison of PPI (%) at 69dB. H) Startle responses (or excessive movement) when no stimuli was present.

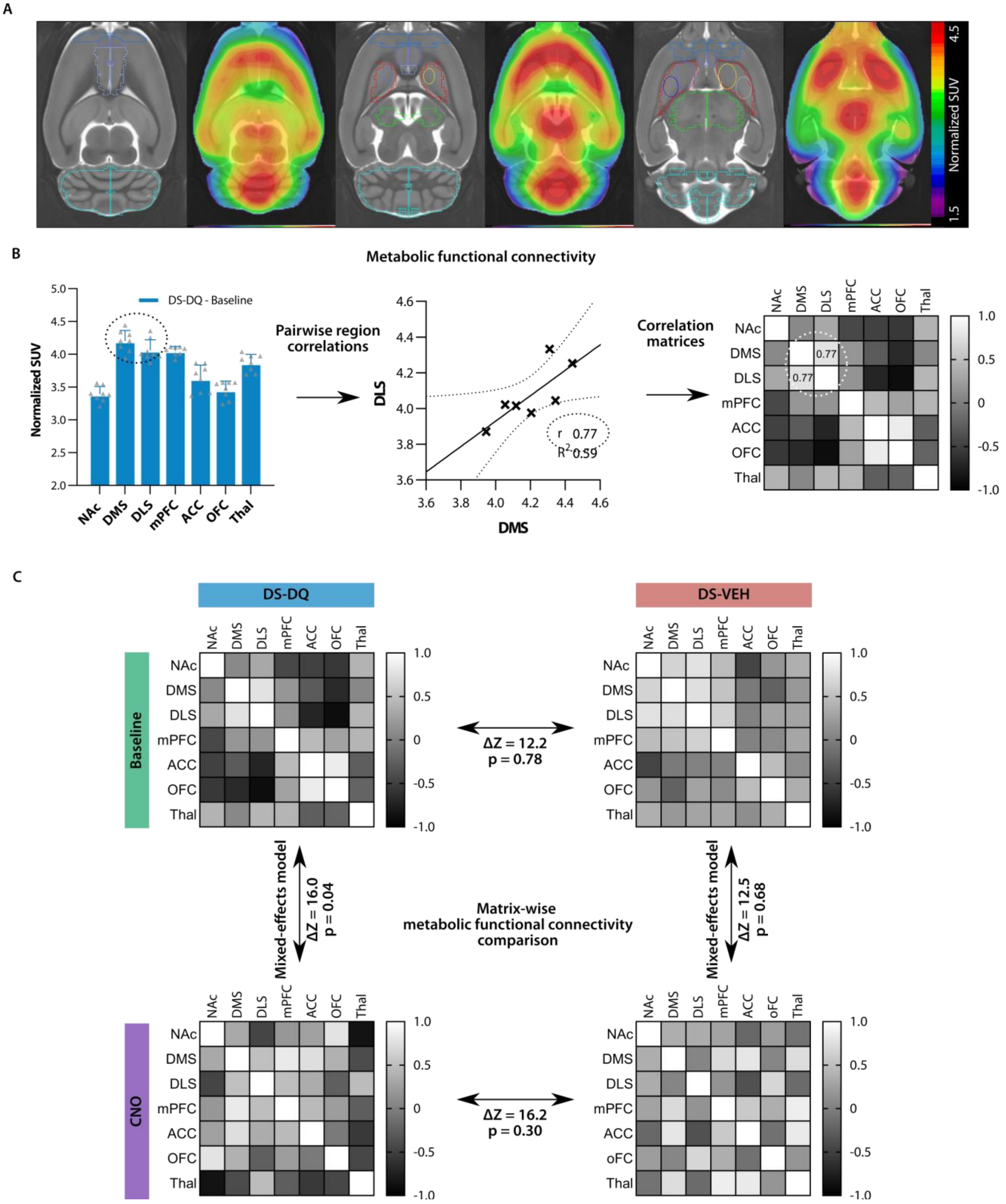


Figure S5: Metabolic functional connectivity. A) Regions drawn on representative structural MR. B) Metabolic functional connectivity explained, pairwise regional connectivity is transformed into four correlation matrices, one for each animal and treatment. C) The four correlation matrices are compared and significant differences are calculated. The functional connectivity within the CSTC circuit is different between baseline and CNO conditions in the DS-DQ animals (p = 0.04).

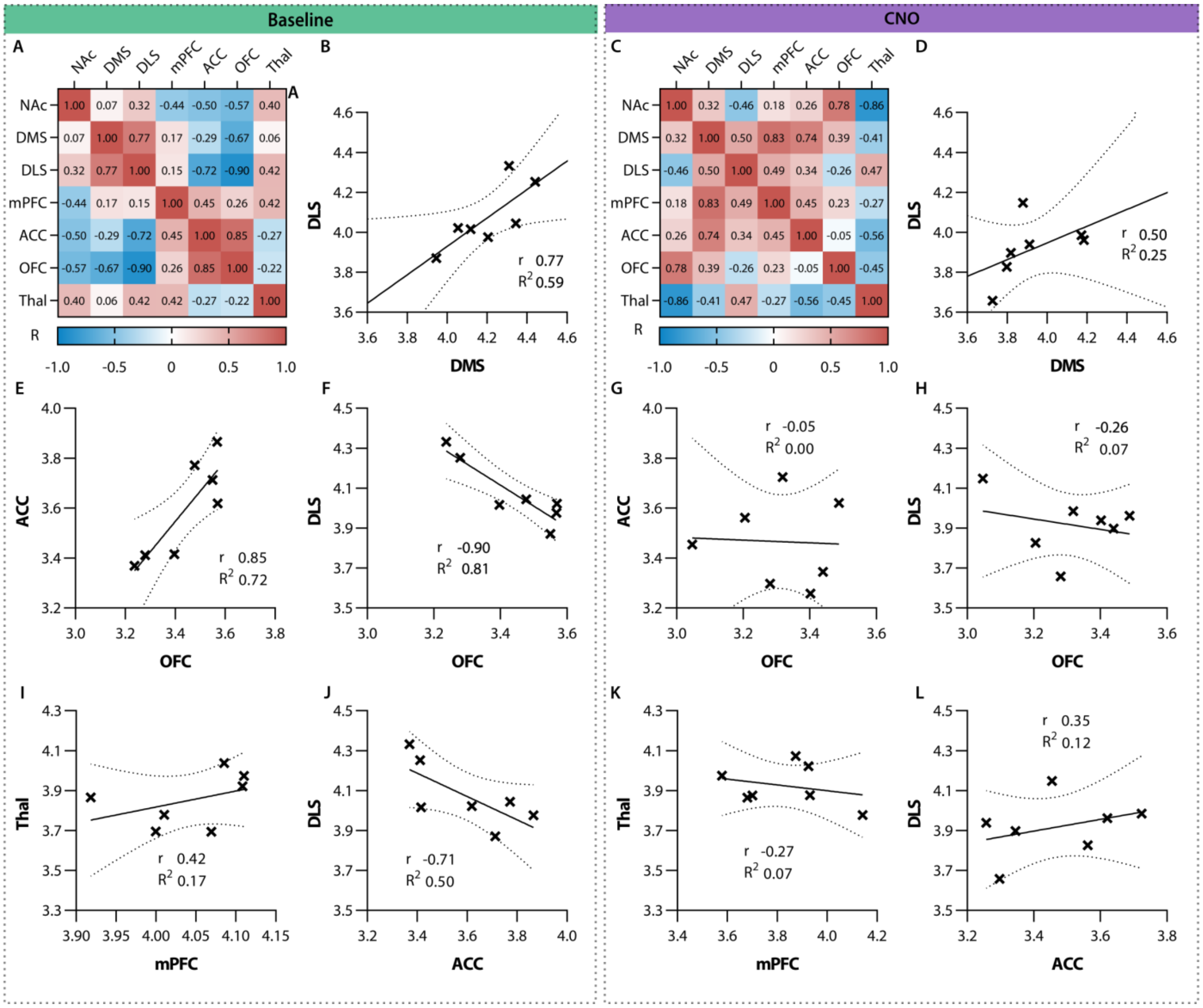


Figure S5: Correlation (metabolic functional connectivity) changes in significant regions at baseline (left side) and after chemogenetic stimulation (right side). A,C) Correlation matrices with Pearson's r values. B,D) DLS / DMS remain correlated after chemogenetic stimulation. E,G) Correlations between ACC / OFC and DLS / OFC are disrupted following chemogenetic stimulation. I,K) Thal / mPFC correlations remain stable. J,L) ACC / DLS correlations are disrupted following chemogenetic stimulation.
